# Lag3 and PD-1 pathways regulate NFAT-dependent TCR signalling programmes during early CD4^+^ T cell activation

**DOI:** 10.1101/2024.06.06.597762

**Authors:** Lozan Sheriff, Alastair Copland, David A.J. Lecky, Reygn Done, Lorna S George, Emma K. Jennings, Sophie Rouvray, Thomas A.E. Elliot, Elizabeth S. Jinks, Lalit Pallan, David Bending

## Abstract

Lag3 and PD-1 are immune checkpoints that regulate T cell responses and are current immunotherapy targets. Yet how they function to control early CD4^+^ T cell activation remains unclear. Here, we show that the PD-1 and Lag3 pathways exhibit layered control of the early CD4^+^ T cell activation process, with the effects of Lag3 more pronounced in the presence of PD-1 pathway co-blockade (CB). RNA-sequencing revealed that CB drove an early NFAT-dependent transcriptional profile, including promotion of ICOS^hi^ T follicular helper (Tfh) cell differentiation. NFAT pathway inhibition abolished CB-induced upregulation of NFAT-dependent co-receptors ICOS and OX40, whilst unaffecting the NFAT-independent gene *Nr4a1*. Mechanistically, Lag3 and PD-1 pathways functioned additively to regulate the duration of T cell receptor (TCR) signals during CD4^+^ T cell re-activation. Our data therefore reveal that PD-1 and Lag3 pathways converge to additively regulate TCR signal duration and NFAT-dependent transcriptional activity during early CD4^+^ T cell re-activation.

**Highlights:**

- PD-1 and Lag3 pathways exhibit layered control of early CD4^+^ T cell activation
- Their co-blockade enhances NFAT-dependent TCR transcriptional programmes
- Inhibition of NFAT signalling reverses the functional effects of PD-1 and Lag3 co-blockade
- Mechanistically, PD-1 and Lag3 function to additively regulate TCR signal duration during re-activation of CD4^+^ T cells

## Introduction

Canonical T cell activation is initiated by the binding of the T cell receptor (TCR) to peptide-loaded major histocompatibility complexes (MHC). The integration of signals from co-stimulatory and co-inhibitory molecules fine-tunes this process^1^ and can alter the threshold for T cell activation^2^. Understanding how T cell activation thresholds are modulated has fundamental implications for the treatment of T cell-mediated diseases and the exploitation of T cell-modulating immunotherapies.

The immune checkpoints programmed cell death protein 1 (PD-1), cytotoxic T lymphocyte antigen 4 (CTLA-4) and lymphocyte activation gene 3 (Lag3), have been identified as critical regulators of T cell activation^3–5^. All three molecules (and for PD-1 also its ligand PD-L1^6^), are current approved targets for cancer immunotherapy^7,8^. Each of these checkpoints is thought to exhibit distinct functions on T cells. PD-1 contains immunoreceptor tyrosine-based inhibitory motifs (ITIMs) that allow it to recruit Src homology-2 domain-containing protein tyrosine phosphatase-2 (SHP-2)^9^ phosphatases to target the TCR^10^ and CD28^11^. CTLA-4 acts on the CD28 pathway, and competes with CD28 for binding to the costimulatory molecules CD80/ CD86^12–14^ and therefore regulates T cell co-stimulation^15^. Lag3, a CD4 structural homologue^16^, binds MHC class II^17^ and is an important regulator of T cell activation and expansion^5^. Indeed, increasing evidence highlights the importance of CD4^+^ T cell tumour surveillance and control through recognition on peptide MHC II complexes^18^. In addition, CD4 T cells are also essential for shaping and enhancing the anti-tumour cytotoxic CD8 response^19,20^.

We recently developed an accelerated adaptive tolerance model that can evaluate the effects of immune checkpoints on CD4^+^ T cell activation *in vivo* ^2,21^. Using this model, we showed that PD-1 acts as a rheostat to control the threshold for T cell activation. Its blockade leads to an enhanced TCR signal strength and the upregulation of co-stimulatory molecules, such as ICOS and OX40^2^. Importantly, we found that the co-inhibitory receptor Lag3 was strongly up-regulated in these tolerogenic CD4 T cells, and so we questioned what effect this pathway would have on T cell function in the context of PD-1 pathway blockade.

To investigate how Lag3 may regulate CD4^+^ T cells in the context of PD-1 pathway blockade, we exploited the Tg4 TCR transgenic model, which is specific for myelin basic protein (MBP) peptide. MBP contains a lysine (K) at position 4, which exhibits unstable MHC binding to the I-A^U^ class II molecule^22,23^. Substituting for a tyrosine (Y) generates an affinity for I-A^U^ which leads to highly stable peptide/MHC II complexes^22,24^ which is potently recognised by the Tg4 transgenic TCR. Given that Lag3 has a preferential function for targeting stable peptide/MHC II complexes^25,26^ we were interested to further understand its role in the context of PD-1/PD-L1 pathway blockade during longer periods of immune activation with [4Y]-MBP.

In this study, we reveal that PD-1 and Lag3 pathways exhibit layered and time-dependent control of CD4^+^ T cell activation. By additively regulating the duration of TCR signalling, the two pathways converge to principally regulate NFAT-dependent distal TCR signalling programmes in CD4^+^ T cells.

## Results

### PD-1 and Lag3 exhibit layered control of early CD4^+^ T cell re-activation

To understand the kinetic relationship between PD-1 and Lag3 expression we analysed a previous RNA-seq time course that investigated the temporal dynamics of gene expression in Tg4 T cells undergoing strong or weak TCR signalling (Figure 1A). These data revealed that the gene encoding PD-1 (*Pdcd1*) was similarly primed by weak or strong TCR signals, behaving largely like an activation marker. In contrast, Lag3 required higher TCR signal strengths and its peak activation was delayed compared to Pdcd1 (Figure 1A and B). In addition, blockade of PD-1 signalling (through anti-PD-L1) led to increased expression of Lag3 on CD4^+^ T cells (Figure 1C). These data led us to hypothesise that Lag-3 and PD-1 may exert layered control of early CD4^+^ T cell re-activation *in vivo*.

**Figure 1:**
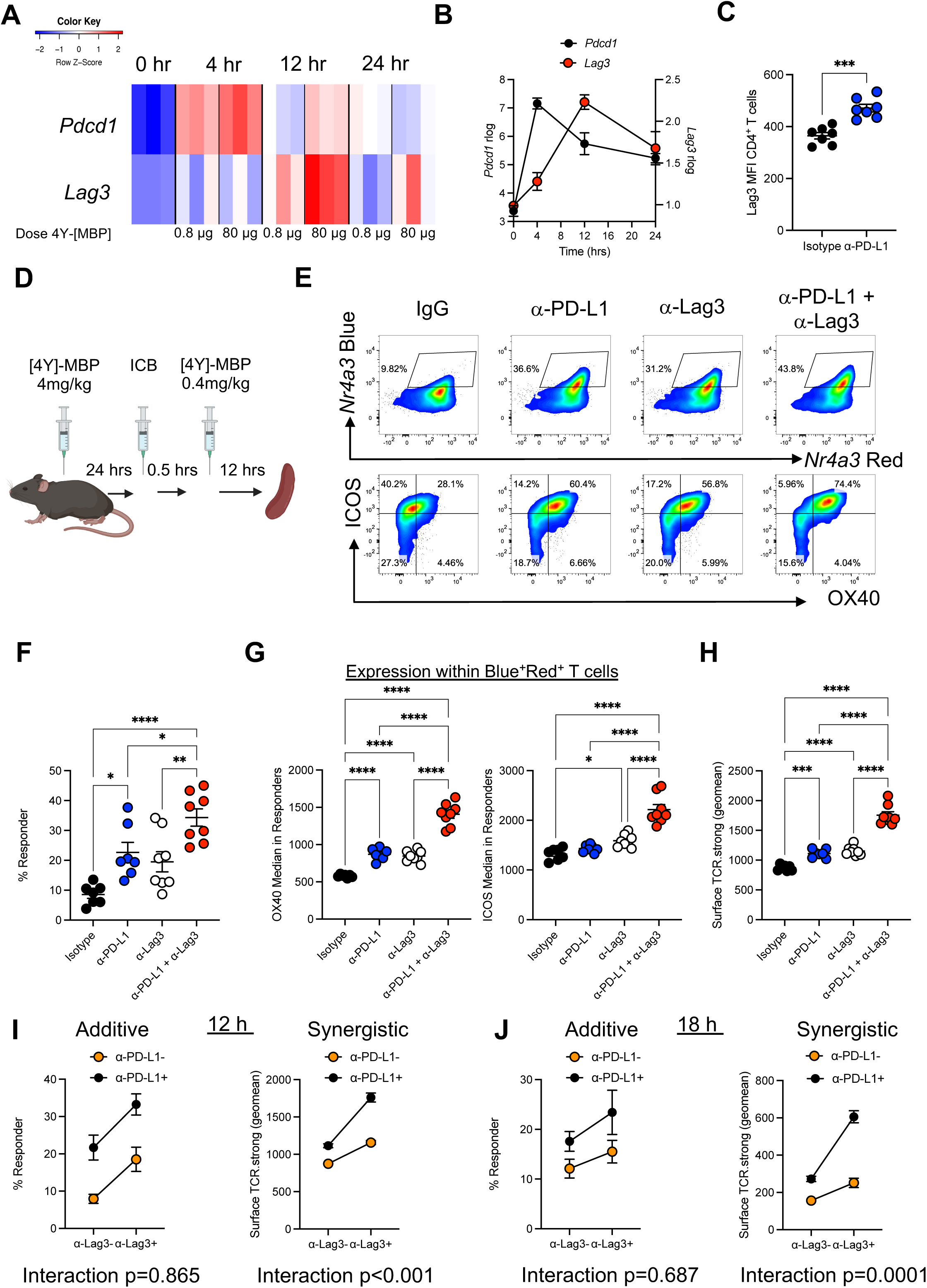
PD-1 and Lag3 exhibit layered control of early CD4^+^ T cell activation. Tg4 TCR transgenic mice were immunised with low (0.8 μg; weak) or high (80 μg; strong) amounts of antigen [4Y]-Myelin basic protein (MBP) peptide s.c. in PBS and then splenic CD4^+^ T cells were analysed by RNA-seq at 0,4,12 and 24hrs post-immunisation for Pdcd1 (PD-1) and Lag3 expression GEO: GSE165817 (**A**). (**B**) Summary of (**A**) for strongly signalled CD4+ T cells. (**C**) Tg4 *Nr4a3*-Tocky *Il10*-GFP mice were immunized s.c. with 4 mg/kg of [4Y] MBP. 24 h later mice were randomized to receive either 0.5 mg isotype or anti-PD-L1 30 min prior to re-challenge with 0.4 mg/kg [4Y] MBP peptide. Splenic CD4^+^ T cells were Lag3 expression 12 h after peptide re-challenge. (**D**) Experimental setup and interpretation for part (**E**). Tg4 *Nr4a3*-Tocky *Il10*-GFP mice were immunized s.c. with 4 mg/kg of [4Y] MBP. 24 h later mice were randomized to receive either 0.5 mg isotype pool (1:1 ratio of rat IgG1 and rat IgG2a), anti-Lag3, or anti-PD-L1 or a combination of both 30 min prior to re-challenge with 0.4 mg/kg [4Y] MBP peptide. Splenic CD4^+^ T cells were analyzed for *Nr4a3*-Blue versus *Nr4a3*-Red analysis at 12 h after peptide re-challenge. Flow plots showing expression of OX40 and ICOS within CD4^+^ T cells. (**F**) Summary data from (B) detailing percentage of responders after 12 h re-challenge. (**G**) and Summary data from (B) detailing MFI (median fluorescence intensity) of OX40 and ICOS within responding CD4^+^ T cells after 12 h re-challenge. (**H**) Summary data detailing geometric mean of ICOS and OX40 (surface TCR.strong) after 12 h re-challenge. (**F**)-(**H**) Isotype (n = 7), anti-Lag3 (n = 7), or anti-PD1 (n =8) or combination therapy (n=8). Bars represent mean ± SEM, dots represent individual mice. Statistical analysis by one-way ANOVA with Tukey’s multiple comparisons test. Two-way anova to test the interaction effects of percentage responders and geometric mean of surface TCR.strong between αLag3 and αPD-L1 at (**I**) 12 h (IgG n=7, anti-PD-L1=7, anti-Lag3=8, and combination n=8) or 18 h (**J**) post re-challenge (IgG n=4, anti-PD-L1=5, anti-Lag3=5, and combination n=5).

Given the potential feedback control of Lag3 by PD-1 pathway blockade we wanted to further understand how Lag3 and PD-1 signalling interact to control CD4^+^ T cell activation. To further investigate the effects of co-blockade (CB) of Lag3 and PD-L1 *in vivo*, we therefore utilised an accelerated adaptive tolerance model^2,21^ to test CB potency effects following T cell reactivation (Figure 1D). In this model, a single high dose of [4Y]-MBP induces transient TCR signalling *in vivo* and upregulation of multiple immune checkpoints, including PD-1 and Lag3, thereby allowing an assessment of their functions^2^. A secondary immunisation subsequently allows a dissection of the roles that Lag3 and PD-L1 play in the activation thresholds for CD4^+^ T cells. CB significantly enhanced Nr4a3-Red^+^Nr4a3-Blue^+^ (i.e. cells that have persistent activity of the NFAT-*Nr4a3* pathway^27^ and henceforth referred to as “responder” cells; mean = 33.3%) compared to isotype control (mean=7.9%) and single agent anti-PD-L1 (mean=21.7%) or anti-Lag3 (mean=18.5%) treatment (Figure 1E and 1F). We have previously identified ICOS and OX40 as markers of PD-1 response in CD4^+^ T cells^2^, so we were interested to further understand their regulation in the context of CB. Analysis of ICOS and OX40 expression within responder cells (to normalise for differences in the frequency of T cell activation *in vivo,* Figure S1) revealed a strikingly large increase in CD4^+^ T cell surface expression of ICOS and OX40 in CB-recipient mice compared to IgG or single agent-treated mice (Figure 1G). We generated a surface protein metric called surface TCR.strong based on the geometric mean of OX40 and ICOS MFI (Figure S1), the transcripts of which we have previously reported to be enriched in melanoma patient responders to Nivolumab^2^. Analysis of surface TCR.strong expression suggested that CB was exerting synergistic effects on the levels of OX40 and ICOS compared to individual treatments (Figure 1H). To further explore the effects of Lag3 and PD-L1 CB on responder and surface TCR.strong metrics, we performed statistical testing to determine whether any interaction (i.e. potential synergy) existed between the treatments (Figure 1I). The effect of CB was additive for the responder population (parallel lines, *p* value for interaction=0.865, ns); however, for the surface TCR.strong metric a significant interaction was detected, highlighting a synergistic effect on ICOS and OX40 expression. As the analysis was performed on only responder populations, this effect is not driven by differential activation states in response to the different treatment regimes. These findings were also detected at a later time point (18hrs) post-secondary stimulation (Figure 1J). These synergistic changes were also detected when ICOS and OX40 were analysed as separate markers (Figure S2). In summary, these data reveal that CB of Lag3 and PD-L1 drives additive increases in the proportion of CD4^+^ T cells that re-activate as well as synergistic increases in markers linked to strong TCR signalling in vivo.

### PD-1 and Lag3 pathway blockade enhances NFAT-dependent TCR transcriptional programmes

To gain an unbiased overview of phenotypic changes in CD4^+^ T cells in response to CB, we repeated the experiments performed in Figure 1D, but this time used FACS to isolate responder cell populations before performing 3’ mRNA sequencing (Figure 2A and 2B). Isolating the responders alone (i.e., Timer^+^ cells) allowed us to normalise for different levels of activation between groups and to home-in on the qualitative changes induced by these treatments. Principal component analysis (PCA) showed that the treatments were largely separated by PC1 (main biological variation), with CB-recipient mice the most distant from isotype and anti-Lag3 and anti-PD-L1-treated mice clustering close together (Figure 2C), with PC2 largely reflecting the batch sorting effects on the cells. Analysis of DEGs revealed that Lag3 and PD-L1 co-regulated genes associated with strong TCR signalling (*Icos, Tnfrsf4, Il21, Tnfrsf9, Maf, Irf8*^2^) but the DEG showing the highest log fold-change was the chemokine receptor *Ccr6* (Table S1, Figure 2D). In addition, there were notable upregulated genes involved in glycolysis/metabolism (*Aldoa, Hk2, Hif1a*), NF-kB regulator gene *Bcl3,* and genes involved in ribosome biogenesis (*Gnl3, Wdr43*). Downregulated genes included the Th2-associated *Gata3,* as well as integrins *Itga4, Itga6* and also genes involved in lymphoid tissue retention (*Sell*, *S1pr1*) (Figure 2E). As we have observed previously^2^ in strongly signalling CD4^+^ T cells, genes encoding molecules involved in TCR signalling were also downregulated (*Cd4, Rasgrp2*). Having noticed a reduction in Gata3 (Th2 master transcription factor), we curated an analysis of genes associated with different T helper subsets (Figure 2F). Visualisation revealed a clear footprint for Tfh cells (notable differences for *Il21, Icos,* and *Maf*) and Th17-associated markers (although *Il17a* and *Il17f* transcripts were not detectable), including *Ccr6*, which was also elevated in anti-PD-L1 treated mice (Figure 2F). Th1-, Th2- and Treg-associated transcripts were less pronounced after CB treatment, albeit with clear variation in expression, reflecting low transcript abundance for many of these genes.

**Figure 2:**
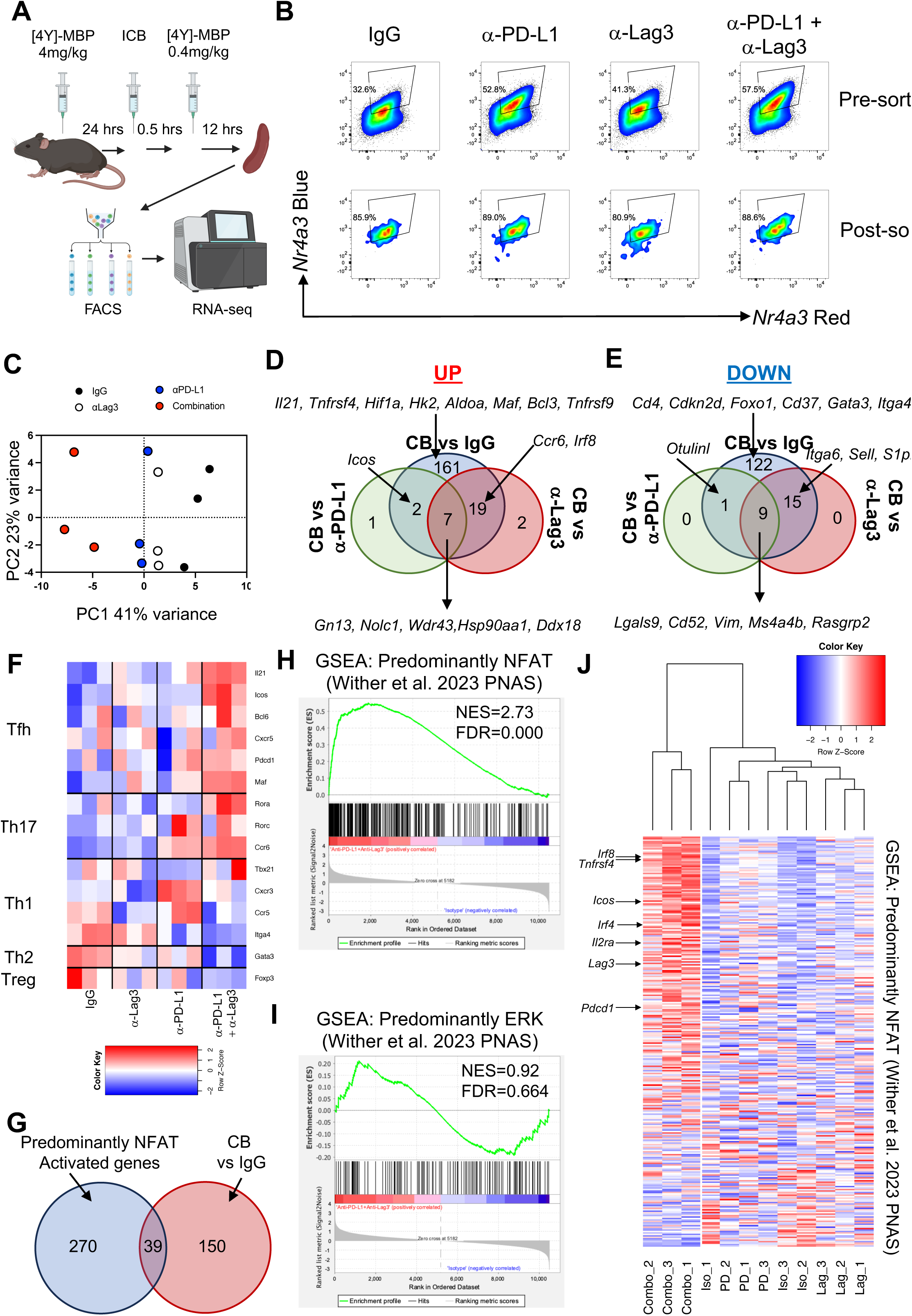
PD-1 and Lag3 blockade enhances NFAT-dependent TCR transcriptional programmes. (**A**) Experimental setup and interpretation for part (B). (**B**) Tg4 *Nr4a3*-Tocky *Il10*-GFP mice were immunized s.c. with 4 mg/kg of [4Y]-MBP. 24 h later mice were randomized to receive either 0.5 mg isotype pool (n=3) (1:1 ratio of rat IgG1 and rat IgG2a), anti-Lag3 (n=3), or anti-PD-L1 (n=3) or combination therapy (n=3) 30 min prior to re-challenge with 0.4 mg/kg [4Y]-MBP peptide. 12 h later splenic CD4^+^ T cell responses were analyzed for *Nr4a3*-Timer Red versus *Nr4a3*-Timer Blue expression in live CD4^+^ Tg4 T cells and sorted for CD4^+^Nr4a3-Blue^+^Nr4a3-Red^+^ T cells by FACS. (**B**) Flow plots showing *Nr4a3*-Timer Blue^+^, *Nr4a3*-Timer Red^+^ pre and post sorting. (**C**) RNA was extracted from the sorted populations and 3′ mRNA sequencing performed. PCA of the normalized expression data identifies 4 clusters. Venn diagram analysis of up (**D**) or down (**E**) regulated DEGs in mice receiving combination therapy compared to the three other experimental conditions. (**F**) Z score heatmap analysis of rLog expression of genes involved in CD4^+^ T helper subsets from Isotype (IgG), anti-Lag3, anti-PD-L1 or CB treated mice. (**G**) Intersection of the 189 genes that were upregulated in CD4^+^ T cell responders between CB treated and Isotype-treated mice with the 309 identified in Wither et al.^29^ as predominantly NFAT dependent. Gene set enrichment analysis (GSEA) of NFAT-(**H**) or ERK- (**I**) dependent signalling based on classification from Wither *et al.*^29^. (**J**) Heatmap visualisation of the normalised counts of the 283/309 genes with largely NFAT-dependent activity visualised across the four treatment conditions and scaled by row values and clustered by columns.

Since transcriptional analysis by RNA-seq provides only a ‘snapshot-in-time’ view, we verified protein levels of the other notable markers identified previously. We performed an identical experiment as in Figure 2A and then evaluated protein intracellular levels of IRF8 (a DEG and part of the TCR.strong metric^2^) and IRF4 (previously linked in this model to strong TCR signalling^2^) (Figure S3A and S3B). Analysis revealed that CB increased levels of IRF8 compared to all three treatment groups, whilst for IRF4 CB increased the levels compared to control and anti-PD-L1 treated mice. In addition, we validated whether the upregulation of *Ccr6* also occurred at the protein level. Analysis of surface CCR6 revealed a sharp, synergistic upregulation (more than 30% of cells now positive for this marker) in response to CB (Figure S3C and S3D) and that CB treated CD4^+^ T cells showed enhanced migration towards the chemokine CCL20 (Figure S3E). In addition we confirmed that the early differentiation of ICOS^hi^ Tfh cells were significantly increased by CB in the Tg4 model (Figure S4A-SD) and that in a polyclonal NP-ova immunisation model, CB also consistently drove increased ICOS^+^ Tfh cell differentiation, albeit with the major contribution coming from PD-1 pathway inhibition (Figure S4E-SH).

Given that *Nr4a3* is a direct NFAT target^27^ and that Tfh cells are known to develop under AP-1 independent NFAT signalling^28^, we wanted to further understand whether the DEGs identified showed any relationship to NFAT signalling pathways. A recent study determined that 1089 genes in T cells are directly regulated by NFAT and ERK pathways during the first 30 hrs^29^ – a similar time frame to our in vivo studies. Through incorporation of inhibitors of NFAT or ERK pathways, Wither *et al.* determined the relative dependence of genes on these signalling axes. We intersected the 189 genes that were upregulated in CD4^+^ T cell responders between CB-treated and isotype-treated mice with the 309 identified in Wither et al. as showing largely NFAT-dependence. Thirty-nine genes were found to overlap between the two data sets (Figure 2G). We performed gene set enrichment analysis (GSEA) based on the annotation of Wither *et al.*^29^ for largely NFAT-dependent (Figure 2H) or largely ERK-driven pathway dependent genes (Figure 2I). GSEA revealed significant enrichment for largely NFAT-dependent genes (Figure 2H, NES=2.73, FDR=0.000) but no enrichment for the predominantly ERK-driven target genes (Figure 2I, NES=0.92, FDR=0.664). Visualisation of the predominantly NFAT-dependent genes across the four conditions showed that only CB-treated CD4^+^ T cells clustered as a discrete group, and hallmark genes such as *Irf8*, *Tnfrsf4*, *Icos*, *Irf4*, *Il2ra*, *Lag3* and *Pdcd1* were all part of the core enrichment set (Figure 2J). Taken together, CB imparts a novel transcriptional signature on CD4^+^ T cells undergoing reactivation *in vivo*, exhibiting hallmarks of NFAT-biased TCR signalling and early promotion of Tfh cell differentiation.

### NFAT pathway inhibition abolishes the major effects of PD-1 and Lag3 pathway co-blockade

Given the significant enrichment of gene and transcriptional programmes that show a large NFAT-dependence, we wanted to functionally explore this pathway. We established a new *in vitro* model to assess the relative contributions of the PD-L1 and Lag3 pathways controlling *in vitro* TCR signal strength in adaptively tolerised T cells (Figure 3A). Splenocytes from Tg4 *Nr4a3*-Tocky *Il10*-GFP mice were isolated 24 hrs after *in vivo* immunisation and cultured in the presence of isotype control or CB in the presence of a dose titration of [4Y]-MBP (Figure 3A to 3C). To demonstrate that this model can reveal manipulations into TCR signalling, we evaluated OX40 and ICOS expression on CD4^+^ T cells. Both OX40 (Figure 3B) and ICOS expression (Figure 3C) were upregulated by CB across a range of agonist peptide concentrations.

**Figure 3:**
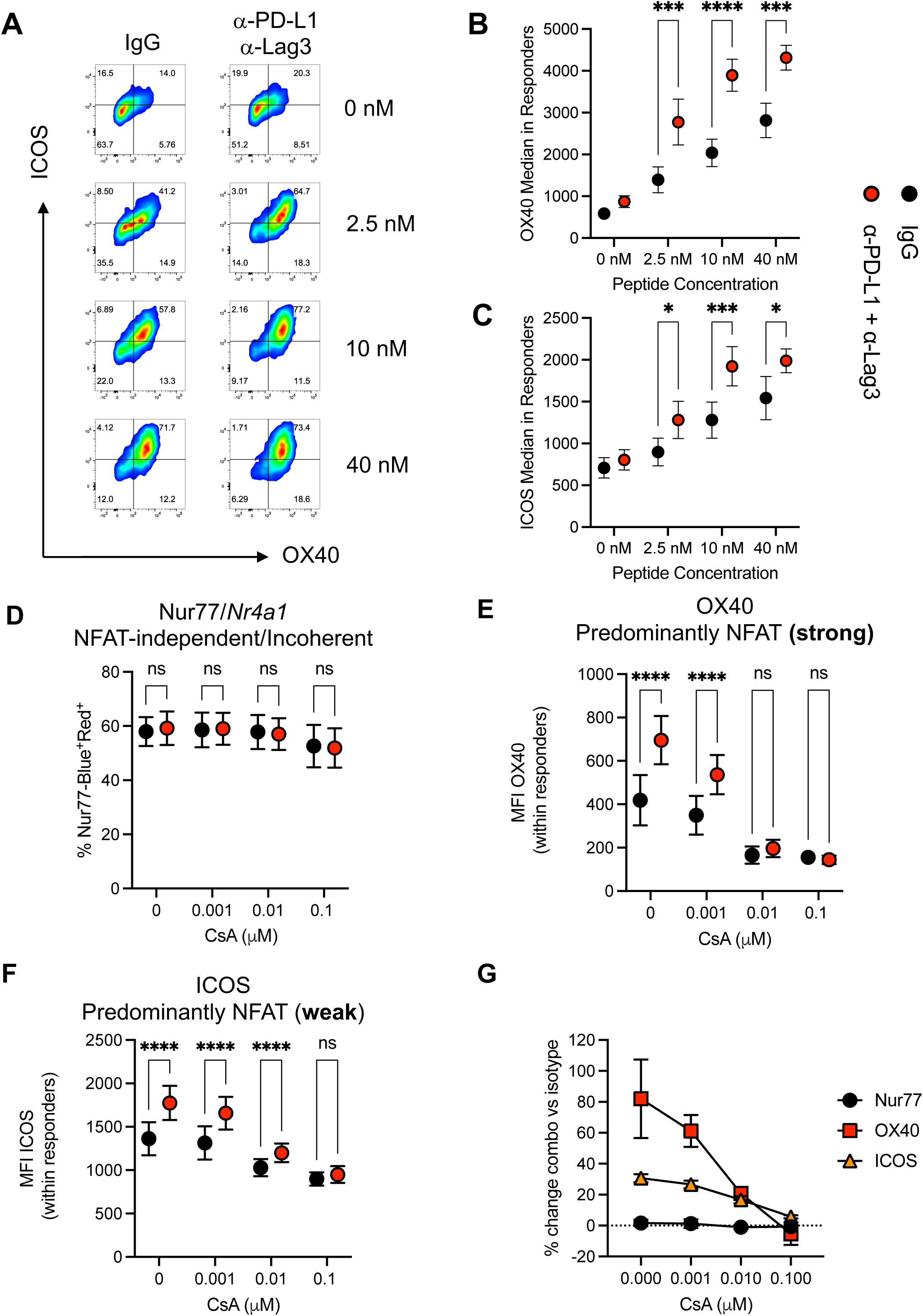
NFAT pathway inhibition abolishes the major effects of PD-1 and Lag3 pathway co-blockade. **(A**) Tg4 *Nr4a3*-Tocky *Il10*-GFP mice were immunized s.c. with 4 mg/kg of [4Y]-MBP peptide. 24 h post immunisation splenocytes were cultured for another 24 h and rechallenged with 0, 2.5, 10, 40 nM [4Y]-MBP peptide in presence of isotype pool or a combination of PD-L1 and αLag-3. Splenic CD4^+^ T cell responses were analyzed for OX40 and ICOS within the responding T cells after 24 h culture. Summary data from (**A**) detailing MFI (median fluorescence intensity) of OX40 (**B**) and ICOS (**C**) within responding CD4^+^ T cells after 24 h culture, (n=7, pooled from 2 independent experiments). (**D-G**) Tg4 Nur77-Tempo *Il10*-GFP mice were immunized s.c. with 4 mg/kg of [4Y]-MBP peptide. 24 h post immunisation splenocytes were cultured for another 24 h and stimulated with 2.5 nM [4Y]-MBP peptide in presence of isotype or a combination of αPD-L1 and αLag-3 and a dose range of cyclosporin A (CsA). After 24hrs the frequency of CD4^+^Nur77-Blue^+^Red^+^ T cells (**D**), median expression of OX40 (**E**), or median expression of ICOS (**F**) in CD4^+^Nur77-Blue^+^Red^+^ T cells was evaluated. (**G**) The percentage change between isotype and combination therapy was calculated for the markers for each dose of CsA. Black circles (isotype treated, n=4) and red circles (combination-treated, n=4, representative of two independent experiments) culture conditions. Bars represent mean ± SEM. Statistical analysis by two-way repeated measures ANOVA with Tukey’s multiple comparisons test.

We then determined the sensitivity of OX40 (predominantly NFAT, strong) and ICOS (predominant NFAT, weak) to NFAT-pathway inhibition (Figure 3D to 3G). We also included Nur77, which we have shown to be NFAT-pathway independent^27^. Nur77 expression was unaffected by CB and the frequency of Nur77-Blue^+^ cells was also unchanged by a range of cyclosporin A (CsA) (Figure 3D). In contrast, CB induced upregulation of OX40 showed enhanced sensitivity to CsA, and at 0.01 µM the increase in OX40 by CB was lost (Figure 3E). CB-induced ICOS upregulation showed a similar pattern to OX40, with the effect of CB lost by 0.1 µM concentration (Figure 3F). Summarising the percentage change revealed that the 3 markers showed alterations in expression by CB which would have been predicted based on the Wither *et al.* model (Figure 3G).

### PD-L1 and Lag3 co-operatively regulate TCR signal duration in CD4^+^ T cells

The maturation of Timer Blue proteins into Timer Red can give an estimation of the signal duration within an actively signalling T cell population (Figure 4A). This is because TCR signalling triggers new formation of Timer Blue proteins, which have a half-life of four hours, before maturation into the longer lived Red proteins from (half-life of 3-5 days). A short TCR signal will thus lead to an increase and then fall in Red proteins, whilst sustained signalling will lead to accumulation of mature Timer Red proteins. We hypothesised that the enhanced NFAT-biased signature observed is a result of CB sustaining TCR signalling for a longer period following reactivation *in vivo* and we therefore re-analysed the levels of Timer Red protein within “responder” cells from Figure 1I and 1J across the four treatment groups. Analysis at 12 hrs post-reactivation in the presence of CB showed a small but significant accumulation of Timer Red proteins compared to the three other groups, however a small increase in Timer Red was also observed in the anti-PD-L1 treated mice (Figure 4B). Analysis at the later 18 hr timepoint revealed that only CB exhibited a significant increase in Timer Red, which demonstrates that these cells have signalled for a longer period (Figure 4C).

**Figure 4:**
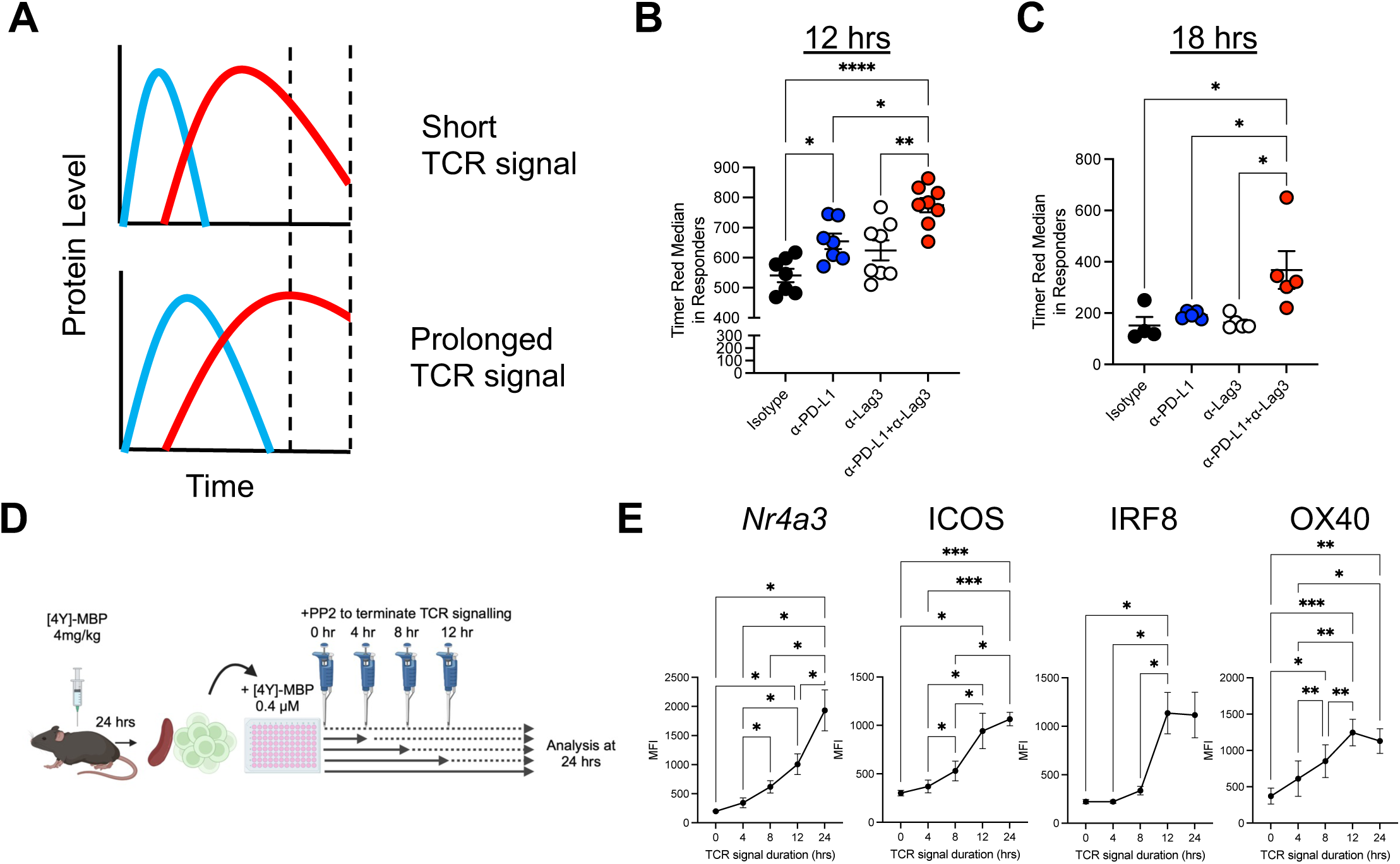
PD-L1 and Lag3 co-operatively regulate TCR signal duration in CD4^+^ T cells. (**A**) Graphical illustration of the levels of Nr4a3-Blue and Nr4a3-Red following a short or prolonged TCR signal. Tg4 *Nr4a3*-Tocky *Il10*-GFP mice were immunized s.c. with 4 mg/kg of [4Y]-MBP. 24 h later mice were randomized to receive either 0.5 mg isotype pool (1:1 ratio of rat IgG1 and rat IgG2a), anti-Lag3, or anti-PD-L1 or CT 30 min prior to re-challenge with 0.4 mg/kg [4Y] MBP peptide. 12 h and 18 h later mice were culled and splenic responses were analysed for *Nr4a3*-Timer Red expression in responder populations. (**B**) Summary of data at 12 h post re-challenge showing red median within responders. Isotype (n = 7), anti-Lag3 (n = 7), or anti-PD1 (n =8) or combination (n=8). Bars represent mean ± SEM, dots represent individual mice. Statistical analysis by one-way ANOVA with Tukey’s multiple comparisons test. (**C**) Summary of data at 18 h post re-challenge showing Timer Red median within responders. Isotype (n =4), anti-Lag3 (n = 5), or anti-PD1 (n =5) or combination (n=5). Bars represent mean ± SEM, dots represent individual mice. (**D**) Experimental setup and interpretation for part (E). Mice were immunized s.c. with 4 mg/kg of [4Y]-MBP (n=6 Nr4a3, ICOS, OX40); n=5, IRF8). 24 h later mice were euthanised, and spleens were harvested. Splenocytes were cultured with 0.1 μM [4Y]-MBP for another 24 h and 10 μM PP2 was added to terminate TCR signalling at different time points (0 h, 4 h, 8 h and 12 h and 24 h) before analysis by flow cytometry. (**E**) Summary data showing MFI (median fluorescence intensity) of *Nr4a3*-Timer Blue, ICOS, OX40 or IRF8 in response to different duration of TCR signalling. Statistical analysis by one-way ANOVA with Tukey’s multiple comparisons test.

We next wanted to test whether key NFAT-dependent markers identified in Figure 2 were regulated by the duration of TCR signalling. We cultured antigen-adapted Tg4 T cells *in vitro* in the presence of [4Y]-MBP and added the Src kinase inhibitor PP2 to terminate TCR signalling after 0, 4, 8, 12 or 24 hours (Figure 4D). Cells remained in culture for the full 24 hours before analysis of Nr4a3-Blue, ICOS, OX40 and IRF8 expression. As expected, Nr4a3-Blue expression continued to increase in parallel with the increase in TCR signal duration (Figure 4E). ICOS expression showed a delayed response but rose prominently from 4 hrs to 12 hrs with a gradual increase up to 24 hrs (Figure 4E). IRF8 showed a similar pattern but its induction required longer periods of TCR signalling (minimum of 12 hrs), and gradually increased thereafter (Figure 4E). OX40 showed a time dependent increase from the initiation of TCR signalling but plateaued from 12 to 24 hrs (Figure 4E). Thus, CB results in an increased duration of TCR signalling, which accounts for the time-dependent and synergistic upregulation of NFAT-dependent markers ICOS, IRF8 and OX40.

## Discussion

Our study demonstrates that the PD-1 and Lag3 pathways exert non-redundant control on the duration of TCR signalling during early CD4^+^ T cell re-activation. PD-1 itself can directly target the TCR^10^ and CD28^11^ molecules, which allows PD-1 to act as a rheostat to control TCR signal strength^2^. PD-1 contains tyrosine-based motifs (ITIM and ITSM^30^) that allow recruitment of SHP-2 phosphatases that mediate its inhibitory function^9^. Lag3 on the other hand has proven more enigmatic in its function. Whilst its binding to MHC Class II is well established^25,26^, Lag3 also has other reported ligands of relevance to T cell biology: galectin-3^31^, fibrinogen-like protein 1 (FLP-1)^32^ and the TCR-CD3 complex^33^. Whilst stable peptide MHC Class II appear more important than FLP-1 for T cell inhibition *in vitro*, the hierarchy of ligands *in vivo* remains unclear^34^. Our findings show that Lag3 works co-operatively with the PD-1/PD-L1 pathway to regulate the T cell activation threshold of CD4^+^ T cells *in vivo*. Mechanistically, co-blockade of PD-L1 and Lag3 results in an increased duration of TCR signalling, which drove synergistic increases in NFAT-dependent pathways.

Interestingly, the effect of Lag3 blockade alone was generally weaker than the PD-L1 pathway *in vivo*, which Vignali and colleagues have also previously noted^33^. In addition, Lag3 had minor effects on the primary CD4^+^ T cell response and its major effects were only exerted in the presence of PD-1 pathway blockade. These findings may be due to broader T cell PD-1 expression and the susceptibility of Lag3 to ADAM-mediated shedding^35^. Indeed, ADAM-resistant forms of Lag3 confer resistance to anti-PD-1 immunotherapy *in vivo*^36^, which is linked to reduced CD4^+^ T cell help for CD8^+^ T cells. It is possible that PD-1 blockade itself may augment the ability of Lag3 to interact with its potential ligands (e.g., MHC class II), which could result in an increased potency of Lag3 in our model, particularly as [4Y]-MBP peptide generates highly stable peptide-MHC II complexes. Given that in the immunised setting blockade of Lag3 resulted in much weaker effects than in the adaptive tolerance conditions, this may reflect the higher levels of Lag3 that are induced on the tolerised CD4^+^ Tg4 T cells.

The fact that PD-1 pathway appears more potent than Lag3 likely also reflects hierarchical levels of T cell tolerance, whereby upon loss of the PD-1 pathway, Lag3 plays a much greater functional role. It is also likely that blockade of the PD-1 pathway may alter the dwell times of T cells on APCs *in vivo*. Tolerised islet antigen-specific T cells showed enhanced motility when scanning DCs in pancreatic draining lymph node^37^. Blockade of PD-L1 or PD-1 led to reduced T cell motility and precipitation of autoimmune diabetes; however blockade of CTLA-4 had no effect on T cell motility^37^ despite past reports that CTLA-4 may limit formation of stable APC:T cell conjugates^38^.

A previous study demonstrated functional synergy of PD-1 and Lag3 in the control of anti-tumour immunity^39^, and this was linked to increases in IFNγ-expressing cells amongst both CD4^+^ and CD8^+^ T cells^39^. Furthermore, several very recent studies have elegantly shown that PD-1 and Lag3 may synergise to promote CD8^+^ T cell exhaustion and hinder their cytotoxicity^40–42^. Unlike CD8^+^ T cells, CD4^+^ T cells have less clearly defined roles in tumour development, exerting anti-tumour effects through effector cytokines, cytotoxicity, and T cell help (reviewed in^43^). However, evidence for pro-tumour development comes from the association of CD4^+^ Foxp3^+^ Tregs with poor clinical outcomes^44^. Nonetheless, expansion of Tfh cells in human tumours has been observed following anti-PD-1 immunotherapy^45^. In solid tumours, Tfh cells likely have a beneficial role^46^ and their tumour infiltration is positively correlated with clinical outcome in human breast cancer^47^. Mechanistically, Tfh cell-derived IL-21 was linked to enhancing CD8^+^ T cell function in mouse models of lung adenocarcinoma^48^. Although this study does did not seek to evaluate the impact of the tumour microenvironment on checkpoint blockade, it has highlighted that re-activation of CD4^+^ T cell under anergising conditions in the presence of CB (i.e., lack of co-stimulation) results in the emergence of NFAT-dependent TCR signalling programmes. In addition, a study assessing cancer-irrelevant immune responses to seasonal flu vaccination describe an increased Tfh cell response in patients on anti-PD-1 therapy^49^. Tfh cell signatures are also positively correlated with immune related adverse events (irAEs) in humans^50^. Interestingly, similar findings have been observed in murine studies using aged mice (18-24 months), which showed the development of IgG-mediated disease that was linked to an IL-21-producing Tfh-like subset^51^. In the present study, combination treatment robustly drove enhanced Tfh cell development, but its most notable effect was the induction of elevated NFAT-pathway dependent molecules such as OX40 and ICOS expression. Given that ICOS is an essential costimulatory molecule for Tfh cell responses *in vivo*^52^, it will be important to understand its wider potential effects on humoral immunity.

The potential manipulation of Tfh cell biology by checkpoint blockade may come with inherent risk. Genetic analysis of SNPs have linked a variant (rs17388568) associated with increased risk of ulcerative colitis^53^ and type 1 diabetes^54^ with an enhanced response to anti-PD-1 in metastatic melanoma patients ^55^. The risk variant maps to a genomic region containing *IL2* and *IL21*, which are critical cytokines involved in regulation of CD4^+^ and Tfh cell responses. Future work should address the extent to which Tfh cell development is altered in patients on immune therapies targeting PD-1 and/ or Lag3 pathways and how these relate to irAEs.

In summary, our study provides new insight into how the PD-1 and Lag3 pathways intersect to govern the duration of TCR signalling during CD4^+^ T cell activation. The co-blockade of these receptors can enhance early NFAT-dependent TCR signalling programmes and the differentiation of Tfh cells.

## Materials and methods

### Mice

*Nr4a3*-Tocky^56^ lines were modified by mating to Tg4 *Il10*-GFP mice as previously described^57^ to generate Tg4 *Nr4a3*-Tocky *Il10*-GFP mice^27^ (unmodified Nr4a3-Tocky lines were originally obtained from Dr Masahiro Ono (Imperial College London) under MTA; the manuscript does not report any findings arising from the use of unmodified *Nr4a3*-Tocky lines or modified *Nr4a3*-Tocky lines that have not been previously reported in the literature. Nur77-Tempo mice^58^ were mated to Tg4 *Il10*-GFP mice to generate Tg4 Nur77-Tempo mice. C57BL/6J mice were purchased from Charles River. All animal experiments were approved by the local animal welfare and ethical review body and authorised under the authority of Home Office licenses P18A892E0A and PP9984349 (held by D.B.). Animals were housed in specific pathogen-free conditions. Both male and female mice typically aged 6-8 weeks were used, and littermates of the same sex were randomly assigned to experimental groups.

### Accelerated adaptive tolerance models and immunotherapy

Tg4 Nur77-Tempo *Il10*-GFP or Tg4 *Nr4a3*-Tocky *Il10*-GFP mice were immunized through subcutaneous injection of 4 mg/kg [4Y] MBP peptide PBS into the flank. For *in vivo* blockade experiments, antibodies were administrated via the intraperitoneal (i.p.) route with 0.5 mg of anti-PD-L1 antibody (clone MIH5 ^59^) or rat IgG2a (clone MAC219, kind gift from Professor Anne Cooke, University of Cambridge), *in vivo* grade anti-Lag3 (clone C9B7W BioLegend) or rat IgG1 (clone MAC221). For combination therapy experiments, anti-PD-L1 and anti-Lag3 or their corresponding isotype were pooled at 1:1 ratios. For re-challenge experiments, 30 minutes after immunotherapy administration a second dose of 0.4 mg/kg [4Y]-MBP peptide in PBS was administered subcutaneously to the contralateral flank. Mice were then euthanised at the indicated time points, and spleens removed to analyse systemic T cell responses.

### Spleen processing and flow cytometry

Splenocytes were dissociated using scissors in 1.2 mL of digestion media containing 1 mg/mL collagenase D (Merck Life Sciences) and 0.1 mg/mL DNase I (Merck Life Sciences) in 1 % FBS (v/v) RPMI for 25 min at 37 °C in a thermo-shaker. Digested tissue was passed through a 70 µM filter (Greiner) and centrifuged at 1500 rpm for 5 min. Red blood cells were lysed in RBC lysis buffer (Invitrogen) for 2 mins on ice before washing in 2% FBS RPMI. Resulting sample was resuspended in FACS buffer (PBS, 2% FBS). Analysis was performed on a BD LSR Fortessa X-20 instrument. The blue form of the Timer protein was detected in the blue (450/40 nm) channel excited off the 405 nm laser. The red form of the Timer protein was detected in the mCherry (610/20 nm) channel excited off the 561 nm laser. A fixable eFluor780-flurescent viability dye (eBioscience) was used for all experiments. Directly conjugated antibodies used in these experiments were CD4 APC or PerCP-Cy5.5 (clone GK1.5, BioLegend), CD4, CD4 Alexa Fluor 700 (clone RM4-4, BioLegend), CD4 BUV737 (clone GK1.5, BD Biosciences),TCR Vβ8.1, 8.2 BUV395 (clone, MR5-2, BD Biosciences) CD4 BUV395 (clone GK1.5, BD Biosciences), CD8a PE-Cy7 (clone 53-6.7, BioLegend), TCRβ FITC (clone H57-597, BioLegend), PD1 PE-Cy7 (clone 29F.1A12, BioLegend), Lag3 APC or PE-Cy7 (clone C9B7W, BioLegend), OX40 APC or PE-Cy7 (clone OX-86, BioLegend), ICOS Alexa Fluor 700 (clone C398.4A, BioLegend), IRF8 PE (clone V3GYWCH, Invitrogen), CD25 PE-Cy7 (clone PC61, BioLegend), CD44 PerCP-Cy5.5. (clone IM7, BioLegend), PD-L1 APC or PE-Cy7 (clone 10F.9G2, BioLegend), IRF4 PE-Cy7 (clone IRF4.3E4, BioLegend), CCR6 APC or PE (clone 29-2L17, BioLegend), CXCR5 APC (clone L138D7, BioLegend), CD19 PE-Cy7 (clone 6D5, BioLegend), CD11b PE-Cy7 (clone M1/70, BioLegend), PD-L1 BV711 (clone 10F.9G2, BioLegend). For intracellular staining of IRF8 or IRF4, the Foxp3 transcription factor staining buffer kit was used according to the manufacturers’ instructions (eBioscience). For cell sorting for RNA-seq experiments, single cell suspensions from the spleens of immunised mice were generated and then multiplexed to sort in batches of four on a FACS ARIA FUSION cell sorter (BD Biosciences). Multiplex sorting was performed using the following CD4 antibodies for the four experimental conditions: Isotype (CD4 APC), anti-PD-L1 (CD4 BUV395), anti-Lag3 (BUV737) and combination therapy (CD4 PerCPcy5.5). A dump channel consisting of CD19 PE-Cy7, CD11b PE-CY7 and CD8-Cy7 was used and cells were sorted as live, CD4^+^CD19^-^CD11b^-^CD8^-^Nr4a3Red^+^Nr4a3Blue^+^ (i.e. “responder” CD4^+^ T cells). For migration assays live, CD4^+^CD19^-^CD11b^-^CD8^-^ were sorted.

### In vitro restimulation of adaptively tolerised CD4^+^ T cells

For the adaptive tolerance model, Tg4 *Nr4a3*-Tocky Tiger (*Il10*-GFP) mice were immunized through subcutaneous injection of 4 mg/kg [4Y] MBP peptide. 24 h later mice were euthanised and spleens were removed and digested as described above. Digested cells were washed once and 5×10^5^ splenocytes cultured for 24 h in 96-well U-bottom plates (Corning) final volume of 200 µl RPMI 1640 containing 10% FCS and penicillin/streptomycin and 55 µM β-mercaptoethanol (Gibco) for the stated time periods in the presence of a range of [4Y]-MBP peptide concentrations. Blocking antibodies (anti-PD-L1 clone MIH5, and anti-Lag3 clone C9B7W) were administered at 20 µg/mL. For functional assessment of the NFAT pathway in promoting OX40 and ICOS expression, Tg4 Nur77-Tempo Tiger (*Il10*-GFP) mice were immunized through subcutaneous injection of 4 mg/kg [4Y] MBP peptide. 24 h later mice were euthanised and spleens were removed and digested as described above. Splenocytes were restimulated with 2.5 nM 4Y [MBP] peptide in the presence of a dose range of cyclosporin a (Cambridge Bioscience) and analysed 24 hrs after *in vitro* culture.

### RNA-seq library preparation and analysis

RNA was extracted from lysates using the PureLink RNA Micro Scale Kit (Invitrogen) according to the manufacturer’s instructions. 5 ng of RNA was used for generation of sequencing libraries using the Quantseq 3’ mRNA-seq Library Preparation kit (Lexogen). Briefly, library generation was commenced with oligodT priming containing the Illumina-specific Read 2 linker sequence. After first strand synthesis, RNA was degraded. Second strand synthesis was initiated by random priming and a DNA polymerase. Random primers contained the illumina-specific Read 1 linker sequence. Double stranded DNA was purified from the reaction using magnetic beads and libraries amplified (18 cycles) and sequences required for cluster generation and sample indexes were introduced. Libraries were normalized and pooled at a concentration of 4 nM for sequencing. Libraries were sequenced using the NextSeq 500 using a High 75 cycle flow cell. Cluster generation and sequencing was then performed and FASTQ files generated. FASTQ files were then processed by Lexogen as follows: Trimming was performed using Cutadapt version 1.18, followed by mapping using STAR version 2.6.1a to the GRCm38 genome. Reads counting as performed using FeatureCounts version 1.6.4. Uniquely mapped read counts conts in the .txt format were used for further analysis using DESeq2 in R version 4.0^60^. A DESeq dataset was created from a matrix of raw read count data. Data were filtered to remove genes with fewer than 30 reads across all samples. Log2 fold change estimates were generated using the DESeq algorithm and shrinkage using the ashr algorithm^61^ to estimate log2 fold changes (lfc). A lfc threshold of 0.2 was set and genes with an adjusted p value<0.05 and an s value<0.005 were used to filter out genes with low expression or negligible gene expression changes. Normalized read counts were transformed using the regularised log (rlog) transformation. Heatmap analysis was performed on the rlog transformed data using the R package gplots. For KEGG pathway analysis clusterProfiler^62^, DOSE^63^ and biomaRt^64^ packages were used. Venn diagrams were generated using the CRAN package VennDiagram. Data are deposited at GEO: GSE260895. Data from Elliot *et al.*^2^, which determined the effect of TCR signal strength on Lag3 and PD-1 expression kinetics, is deposited at GEO.

### Gene set enrichment analysis (GSEA)

Normalised counts for combination-treated or isotype-treated CD4 T cells from RNA-seq analysis performed in DESeq2 were fed in to GSEA 4.3.3^65,66^. 1000 permutations were performed using the gene_set permutation using NFAT and ERK-dependent gene sets as annotated in Wither *et al.*^29^. For the predominantly NFAT-dependent gene set, 283 out of 309 were detected in our dataset. For the predominantly ERK-dependent gene set 124 out of 136 were detected in our dataset.

### TCR signal duration termination by PP2 administration

*Nr4a3*-Tocky Tg4 Tiger mice were subcutaneously injected with 4 mg/kg [4Y] MBP peptide. Mice were culled 24 h later and spleens were dissociated and single cell suspensions of splenocytes were made using the digest method as described above. Cells were washed once and cultured at 5×10^5^ cells per well on 96-well U-bottom plates (Corning) in presence of 0.1 μM [4Y]-MBP peptide in a final volume of 200 μl RPMI1640 + L-glutamine (GIBCO) containing 10% FCS and 1% penicillin/streptomycin (Life Technologies). Inhibitors were dissolved in DMSO. PP2 (Sigma, 10 μM) was added at different time points. Cells were incubated at 37°C and 5% CO2 and analysed at the indicated time points on flow cytometry.

### Immunization for Tfh cell responses

C57BL/6J mice were immunized with alum-precipitated (9% aluminum potassium sulfate (Sigma-Aldrich)) NP-OVA conjugate (NP-conjugated ovalbumin), alum NP-OVA. These were respectively prepared by mixing endotoxin-free NP-conjugated ova (kind gift from Prof Kai Toellner, University of Birmingham) with a 9% alum solution. Per mouse a total of 20 μg NP-OVA was mixed with same volume of 9% alum and pH was adjusted using NaOH and HCL. After washing, 20μg NP-ova/Alum was resuspended in a total volume of 20μl PBS and injected into the left footpad. 24 h later mice were administrated via the intraperitoneal (i.p.) route with 0.5 mg anti-PD-L1 (clone MIH5), anti-Lag3 (clone C9B7W), combination therapy of their respective isotype control (pooled at 1:1 ratios). On day 8, mice were euthanised and the popliteal lymph node was harvested. LNs were forced through 70 µM filters, before cells were stained for flow cytometric analysis.

### Migration assays

To measure migration, FACS purified CD4^+^ T cells were cultured in a 5 μm polycarbonate 24-well transwell system (Corning). T cells were loaded into the top chamber at a density of 2.5 x 10^5^ cells in 0.1 mL complete media, with the bottom chamber containing either 0 or 100 ng/mL CCL20 (BioLegend) in 0.6 mL complete media. After 4 hours, the contents of the bottom chamber were harvested and washed extensively with PBS (+ 2 mM EDTA and 2% FCS), followed by enumeration by flow cytometry. AccuCheck couning beads (Invitrogen) were used to calculate the absolute number of CD4^+^ T cells in each sample. Migration index was calculated by normalising cells in the lower chamber compared to the medium alone control from isotype-treated mice.

### Statistical Analysis

For non-RNA-sequencing analysis, statistical analysis was performed on Prism 10 (GraphPad) software. Flow cytometry data were analyzed using FlowJo software (BD Biosciences). Statistical analysis for comparison of more than two means was performed using One-Way ANOVA with Tukey’s multiple comparisons test. For analysis of two factor variable experiments, a Two-way repeated measures ANOVA was performed with Tukey’s multiple comparisons tests. For detection of synergy, a Two-way ANOVA test was performed factoring in the variables anti-PD-L1 and anti-Lag3 to test for significance of interaction. For non-parametric data a Kruskal Wallis test was performed with Dunn’s multiple comparisons. Variance is reported as mean ± SEM unless stated otherwise; data points represent individual mice. ^∗^p = < 0.05, ^∗∗^p = < 0.01, ^∗∗∗^p = < 0.001, ^∗∗∗∗^p = < 0.0001.

## Acknowledgements and funding sources

Work is funded by an MRC Career Development award (MR/V009052/1 to D.B.) and a Lister Institute of Preventive Medicine Fellowship (to D.B.). We thank Genomics Birmingham for performing RNA-seq library preparation and sequencing. We thank Ferdus Sheik and Dr Guillaume Desanti in the Birmingham Flow cytometry facility for support with cell sorting. Diagrams in the figures were generated using <u>BioRender.com</u>.

## Supplementary Figures and Tables

**Figure S1. Gating strategy and generation of surface TCR.strong metric (related to Figure 1)**

**Figure S2. Synergistic upregulation of ICOS and OX40 in response to Lag3 and PD-L1 co-blockade (related to Figure 1)** Data from Figure 1I and 1J for the compound surface TCR.strong measurement were split and displayed as the individual components OX40 and ICOS. Top details 12 hr time point, bottom 18 hrs. Bars represent mean ± SEM, statistical analysis by two-way Anova.

**Figure S3. Validation of RNA-seq analysis at protein level (related to Figure 2)** Tg4 *Nr4a3*-Tocky *Il10*-GFP mice were immunized s.c. with 4 mg/kg of [4Y]-MBP. 24 h later mice were randomized to receive either 0.5 mg isotype pool (1:1 ratio of rat IgG1 and rat IgG2a), anti-Lag3, or anti-PD-L1 or a combination therapy 30 min prior to re-challenge with 0.4 mg/kg [4Y]-MBP peptide. 12 h later mice were euthanised and splenic responses were analysed by flow cytometry. (**A**) Flow cytometry plots showing expression of IRF8 and IRF4 in total CD4^+^ T cells and (**B**) summary data showing median expression levels of IRF8 and IRF4. (**C**) Flow cytometry plots showing CCR6 expression (gated on responder T cells) and (**D**) Two-way anova to test the interaction for the treatments in driving expression of CCR6. (**A**)-(**D**) Isotype (n=5), anti-Lag3 (n=5), or anti-PD1 (n=5) or CB treatment (n=6). Bars represent mean ± SEM, dots represent individual mice. Statistical analysis by one-way ANOVA with Tukey’s multiple comparisons test. (**E**) CD4^+^ T cells were FACS purified and cultured in transwells with either 0 or 100 ng/mL of CCL20. 4 hours later the migration index was calculated for the four experimental conditions (Isotype (n=6), anti-Lag3 (n=8), or anti-PD1 (n=8) or CT (n=8), pooled from two independent experiments.

**Figure S4: PD-1 and Lag3 blockade enhances ICOS^hi^ Tfh cell differentiation (related to Figure 2)** (**A**) Experimental setup and interpretation for part (**B**). Tg4 Nur77-Tempo *Il10*-GFP mice were immunized s.c. with 4 mg/kg of [4Y]-MBP. 24 h later mice were randomized to receive either 0.5 mg isotype pool (1:1 ratio of rat IgG1 and rat IgG2a), anti-Lag3, or anti-PD-L1 or CT 30 min prior to re-challenge with 0.4 mg/kg [4Y]-MBP peptide. 18 h later mice were euthanised and splenic responses were analysed for PD-1, CXCR5 and ICOS expression. (**C**) Frequency of Tfh cells amongst the CD4^+^ T cells, (**D**) median ICOS expression in Tfh cells. Isotype (n=4), anti-Lag3 (n =5), anti-PD1 (n =5) or CT (n=5). (**E**) Experimental setup and interpretation for part (**F-H**). (**F**) Mice were immunized with 20 μg NP-OVA/ alum in a total volume of 20 μl subcutaneously into the left foot pad. 24 h later mice were injected randomly either with 0.5 mg isotype pool (1:1 ratio of rat IgG1 and rat IgG2a), anti-Lag3, or anti-PD-L1 or combination treatment. At day 8 post immunization mice were sacrificed and popliteal lymph nodes were harvested for analysis by flow cytometry. Flow cytometry plots showing expression of PD-1 versus CXCR5 or CD4 versus ICOS in live CD4^+^ T cells. (**G-H**) summary data detailing the percentage of Tfh cells **(G**), ICOS median in CD4^+^ T cells (**H**). Isotype (n=14), anti-Lag3 (n = 14), anti-PD1 (n =14) or combination therapy (n=13), data are pooled from two independent experiments. Bars represent mean ± SEM (**C, D** and **H**) or median (**G**), dots represent individual mice. Statistical analysis by one-way ANOVA with Tukey’s multiple comparisons test (**C, D**, **H**) or Kruskal Wallis test with Dunn’s multiple comparisons test (**G**).

**Table S1: DEGs between Isotype, anti-PD-L1 and anti-Lag3 and CB treated groups (related to Figure 2)**

