## Supplementary Figures 1 to 4 for "Lag3 and PD-1 pathways regulate NFAT-dependent TCR signalling programmes during early CD4^+^ T cell activation"

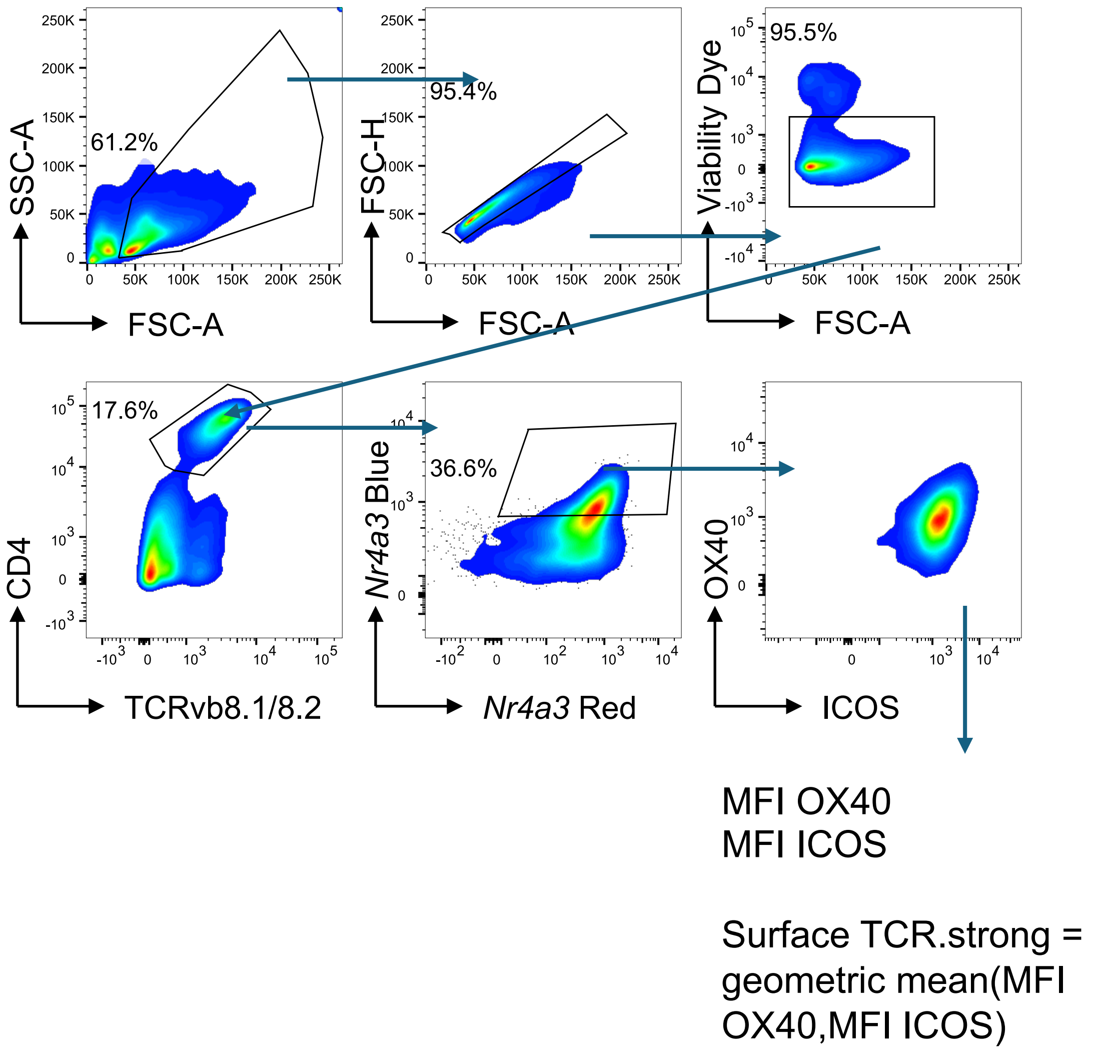


**Figure S1. Gating strategy and generation of surface TCR.strong metric (related to Figure 1)**

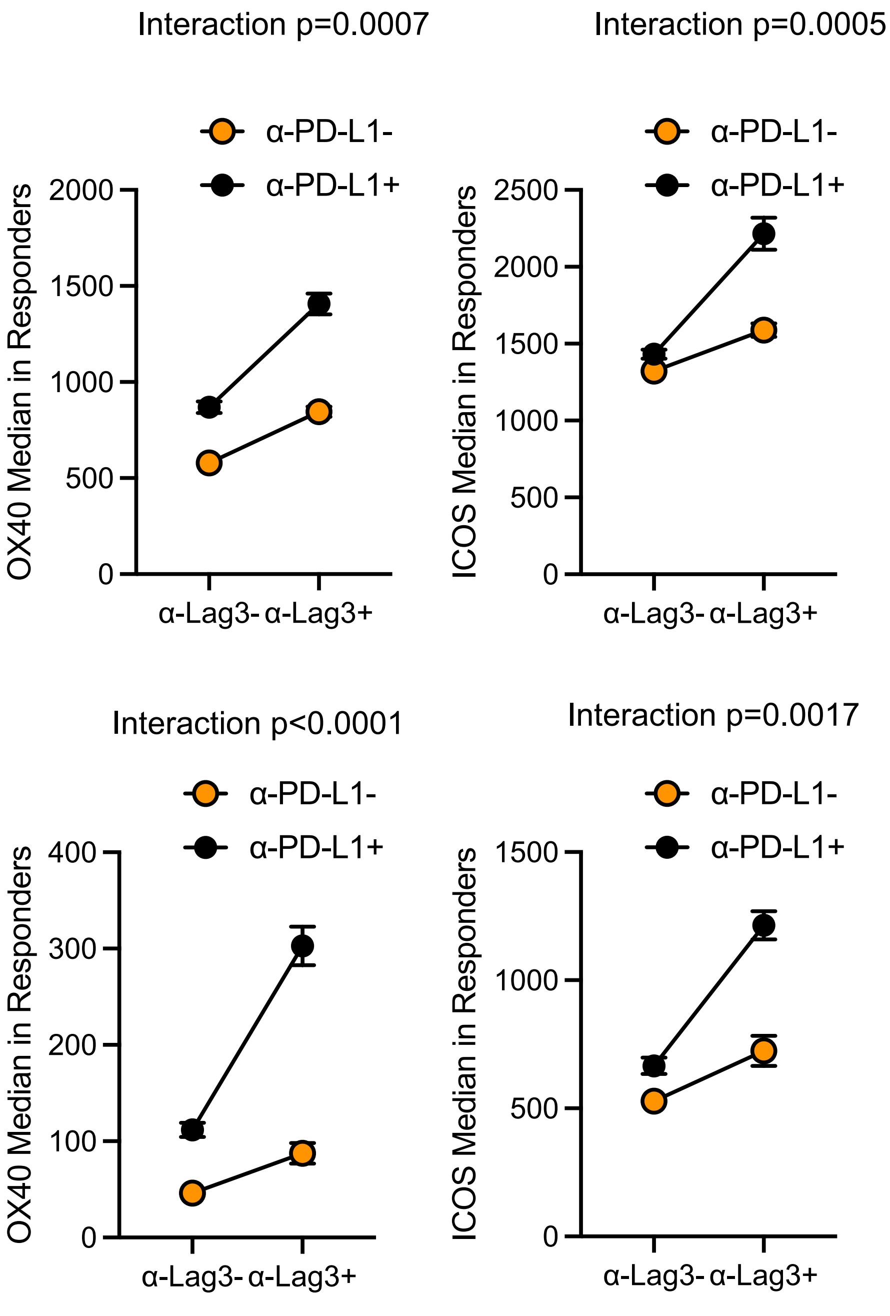


**12 hrs post CN**

**18 hrs post CB**

**Figure S2. Synergistic upregulation of ICOS and OX40 in response to Lag3 and PD-L1 co-blockade (related to Figure 1)**

Data from Figure 1I and 1J for the compound surface TCR.strong measurement were split and displayed as the individual components OX40 and ICOS. Top details 12 hr time point, bottom 18 hrs. Bars represent mean ± SEM, statistical analysis by two-way Anova.

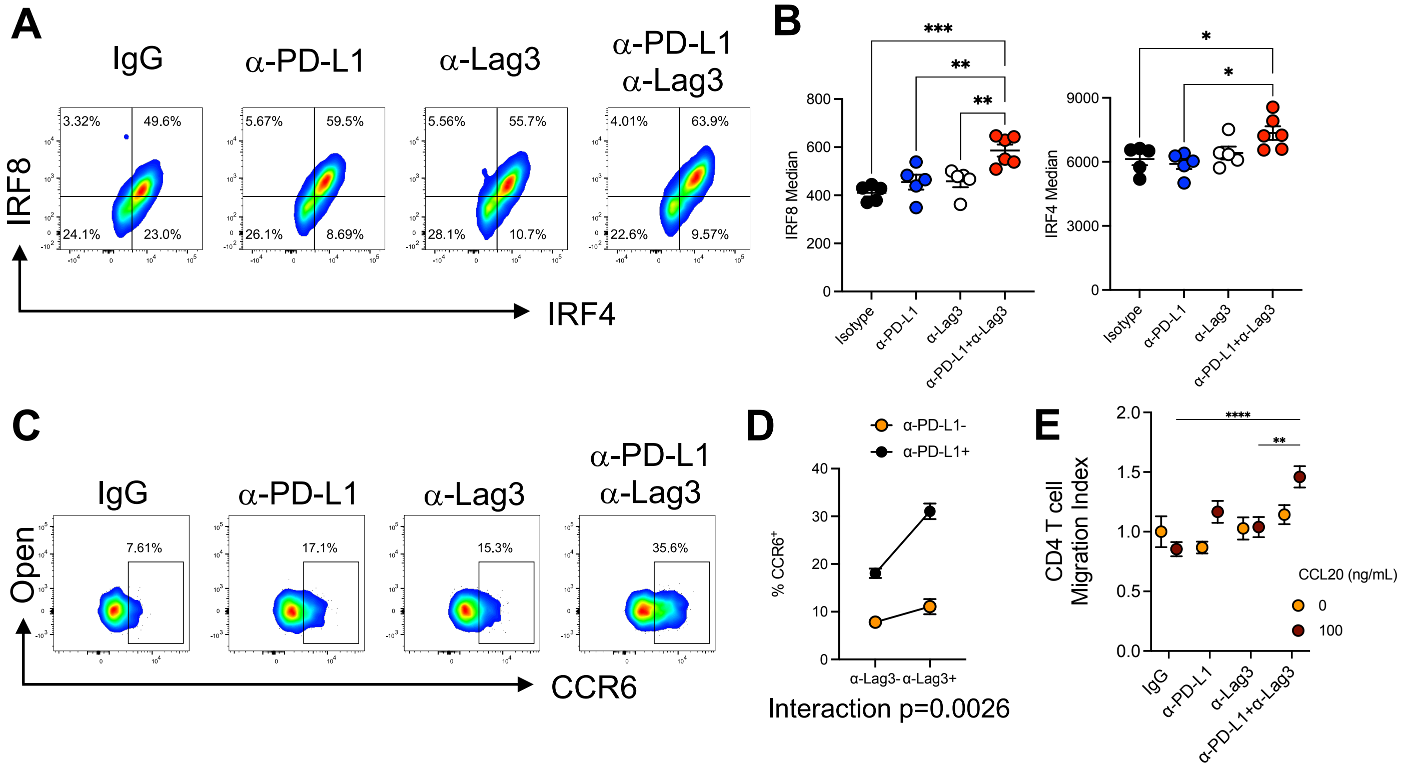


**Figure S3. Validation of RNA-seq analysis at protein level (related to Figure 2)**

Tg4 *Nr4a3*-Tocky *Il10*-GFP mice were immunized s.c. with 4 mg/kg of [4Y]-MBP. 24 h later mice were randomized to receive either 0.5 mg isotype pool (1:1 ratio of rat IgG1 and rat IgG2a), anti-Lag3, or anti-PD-L1 or a combination therapy 30 min prior to re-challenge with 0.4 mg/kg [4Y]-MBP peptide. 12 h later mice were euthanised and splenic responses were analysed by flow cytometry. (**A**) Flow cytometry plots showing expression of IRF8 and IRF4 in total CD4$^{+}$ T cells and (**B**) summary data showing median expression levels of IRF8 and IRF4. (**C**) Flow cytometry plots showing CCR6 expression (gated on responder T cells) and (**D**) Two-way anova to test the interaction for the treatments in driving expression of CCR6. (**A**)-(**D**) Isotype (n=5), anti-Lag3 (n=5), or anti-PD1 (n=5) or CB treatment (n=6). Bars represent mean ± SEM, dots represent individual mice. Statistical analysis by one-way ANOVA with Tukey's multiple comparisons test. (**E**) CD4$^{+}$ T cells were FACS purified and cultured in transwells with either 0 or 100 ng/mL of CCL20. 4 hours later the migration index was calculated for the four experimental conditions (Isotype (n=6), anti-Lag3 (n=8), or anti-PD1 (n=8) or CT (n=8), pooled from two independent experiments.

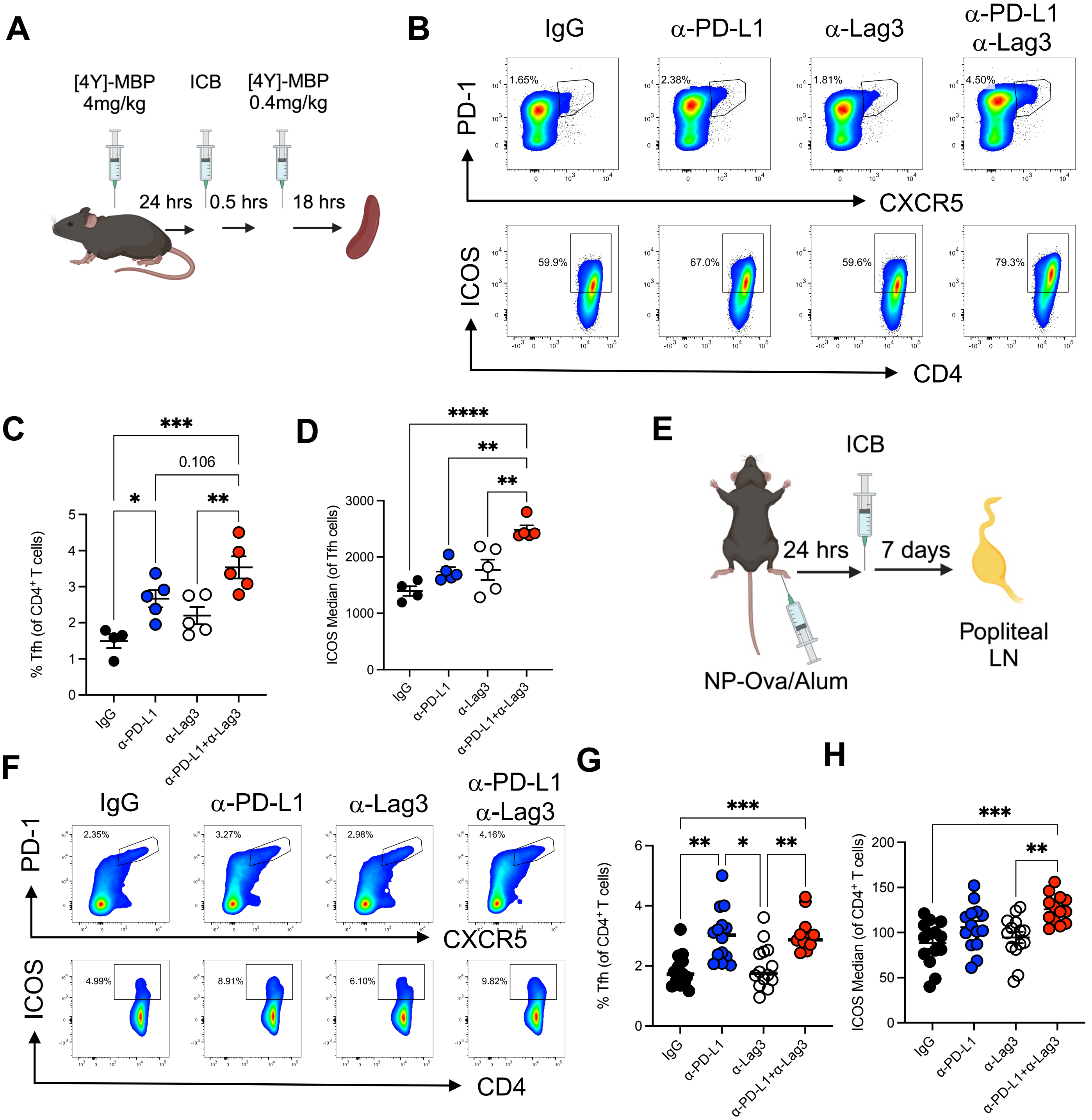


**Figure S4: PD-1 and Lag3 blockade enhances ICOS$^{hi}$ Tfh cell differentiation (related to Figure 2)**

(**A**) Experimental setup and interpretation for part (**B**). Tg4 Nur77-Tempo *Il10*-GFP mice were immunized s.c. with 4 mg/kg of [4Y]-MBP. 24 h later mice were randomized to receive either 0.5 mg isotype pool (1:1 ratio of rat IgG1 and rat IgG2a), anti-Lag3, or anti-PD-L1 or CT 30 min prior to re-challenge with 0.4 mg/kg [4Y]-MBP peptide. 18 h later mice were euthanised and splenic responses were analysed for PD-1, CXCR5 and ICOS expression. (**C**) Frequency of Tfh cells amongst the CD4$^{+}$ T cells, (**D**) median ICOS expression in Tfh cells. Isotype (n=4), anti-Lag3 (n =5), anti-PD1 (n =5) or CT (n=5). (**E**) Experimental setup and interpretation for part (**F-H**). (**F**) Mice were immunized with 20 μg NP-OVA/ alum in a total volume of 20 μl subcutaneously into the left foot pad. 24 h later mice were injected randomly either with 0.5 mg isotype pool (1:1 ratio of rat IgG1 and rat IgG2a), anti-Lag3, or anti-PD-L1 or combination treatment. At day 8 post immunization mice were euthanised and popliteal lymph nodes were harvested for analysis by flow cytometry. Flow cytometry plots showing expression of PD-1 versus CXCR5 or CD4 versus ICOS in live CD4$^{+}$ T cells. (**G-H**) summary data detailing the percentage of Tfh cells (**G**), ICOS median in CD4$^{+}$ T cells (**H**). Isotype (n=14), anti-Lag3 (n = 14), anti-PD1 (n =14) or combination therapy (n=13), data are pooled from two independent experiments. Bars represent mean ± SEM (**C, D** and **H**) or median (**G**), dots represent individual mice. Statistical analysis by one-way ANOVA with Tukey's multiple comparisons test (**C, D, H**) or Kruskal Wallis test with Dunn's multiple comparisons test (**G**).
